# TREM2-dependent senescent microglia conserved in aging and Alzheimer’s disease

**DOI:** 10.1101/2023.03.20.533401

**Authors:** N Rachmian, S. Medina, U. Cherqui, H. Akiva, D Deitch, D Edilbi, T. Croese, TM. Salame, JM. Peralta Ramos, L. Cahalon, V. Krizhanovsky, M. Schwartz

## Abstract

Dementia in general, and Alzheimer’s disease (AD) in particular, are age-related diseases^1,2^. AD is associated with multiple causative factors^3,4^, among which local brain inflammation plays a significant role^5^. Microglia, the brain-resident immune cells^6,7^, are activated along the disease course^7^. Yet, their contribution to the disease progression is still controversial. Here, using high-throughput mass cytometry for microglial immuno-phenotyping, we identified accumulation of senescent microglia in several pathologies associated with cognitive decline. These senescent microglia have a unique profile conserved across the multiple conditions investigated, including aging, mouse models of amyloidosis, and tauopathy. Moreover, we found that the expression of markers of senescence correlates with levels of TREM2, whose polymorphism was identified by GWAS as an AD risk factor^8,9^. A TREM2-null AD mouse model showed lower levels of senescent microglia, relative to TREM2-intact AD mice. Senolysis using the drug ABT-737^10,11^ in an AD mouse model reduced the abundance of TREM2-senescent microglia without affecting levels of TREM2-dependent activated microglia, ameliorated cognitive deficits, and reduced brain inflammation. These results reveal the unexpected contribution of TREM2 to accumulation of senescent microglia in AD pathology, an effect that must be considered when targeting TREM2 as a therapeutic approach.

## Main

Alzheimer’s disease (AD) is a chronic neurodegenerative disease characterized by several disease-escalating factors^3,4^, among which local brain inflammation, mediated by innate immune cells, significantly impacts cognitive loss^5^. Genome-wide association studies (GWAS) have identified immune-related genes as risk factors in disease onset and severity, some of which are expressed by microglia^8,9,12^, the resident innate immune cells of the brain^6,7^. Under pathological conditions, the microglia undergo changes in their fate^13^. In animal models of amyloidosis, these changers include Triggering receptor expressed on myeloid cells 2 (TREM2)-dependent activation^13-16^. TREM2 polymorphism is considered a risk factor in late-onset AD, identified by GWAS^8,9^. In animal models of AD, TREM2-expressing activated microglia were found surrounding amyloid plaques^17^. Deficiency in TREM2 in an animal model was shown to be associated with elevated levels of amyloid plaques, with no effect on the soluble amyloid beta fraction^18^. Moreover, TREM2 facilitates microglial proliferation and their acquisition of the previously described disease-associated microglia (DAM) profile^13,17^.

In addition to genetic factors, aging has been recognized as a major risk factor for many diseases, including AD^1,2^. Several groups have attributed to the immune system a key role in supporting brain maintenance and functional plasticity^19,20,21^ and linked AD to aging of the immune system^22,23^. One of the main hallmarks of the aging process is the accumulation of senescent cells^24,25^. Senescent cells are defined by persistent cell cycle arrest, consistent with a DNA damage response, and a Senescence-Associated-Secretory-Phenotype (SASP)^25^. Through their continuous secretion of pro-inflammatory cytokines, senescent cells contribute to the chronic inflammation associated with aging^25^. This highlights the potentially significant role of senescent cells in neurodegenerative conditions. Key questions are whether among the senescent cells within the diseased or aged brains are microglia, and if so, whether activated and senescent microglia represent a continuum of two cell states, or outcomes of distinct cellular pathways, and whether selectively removal of senescent microglia, will have a positive effect on cognitive performance.

## Signature of senescent microglia

To identify and characterize senescent microglia, we screened for expression of a panel of unique markers for brain myeloid cells and of senescence (Extended Data Table 1). Using mass cytometry (CyTOF), we observed a distinct population of microglia with a characteristic senescent signature in the brains of aged mice (24 months) (Fig. 1a-c), mouse models of familial AD (5xFAD^26^) (Fig. 1d-f), and tauopathy (DMhTAU^27^) (Fig. 1g,h). Microglia were defined as myeloid cells that do not express the monocyte markers Ly-6c and CCR2. Resting microglia were defined as cells that did not express the activation marker CD11c, but expressed classic microglial markers (CD11b, P2RY12, CX3CR1, TMEM119). Senescent microglia were defined as cells expressing senescence hallmarks, including different components of the senescent phenotype (p16, p19, pp38, γH2AX, p21, p53). Interestingly, however, the senescent microglia also expressed characteristic markers of homeostatic microglia, including TMEM119, P2RY12, CX3CR1, and some markers of activity, such as TREM2, CD38, ApoE, C5aR, Cd115, SiglecH and CD39 (Fig 1a,d,g). We detected a higher percentage of senescent microglia in the aged group compared to the adult mice (Fig. 1c). In addition, we found an accumulation of senescent microglia in 5xFAD mice relative to age-matched wild-type (WT) mice (Fig. 1f). The signature of the senescent microglia was conserved across the different models, and included TREM2 expression (Fig. 1g,h). The conserved signature of the senescent microglia accumulated in aging and AD, could explain the strong association between these conditions.

**Fig. 1.**
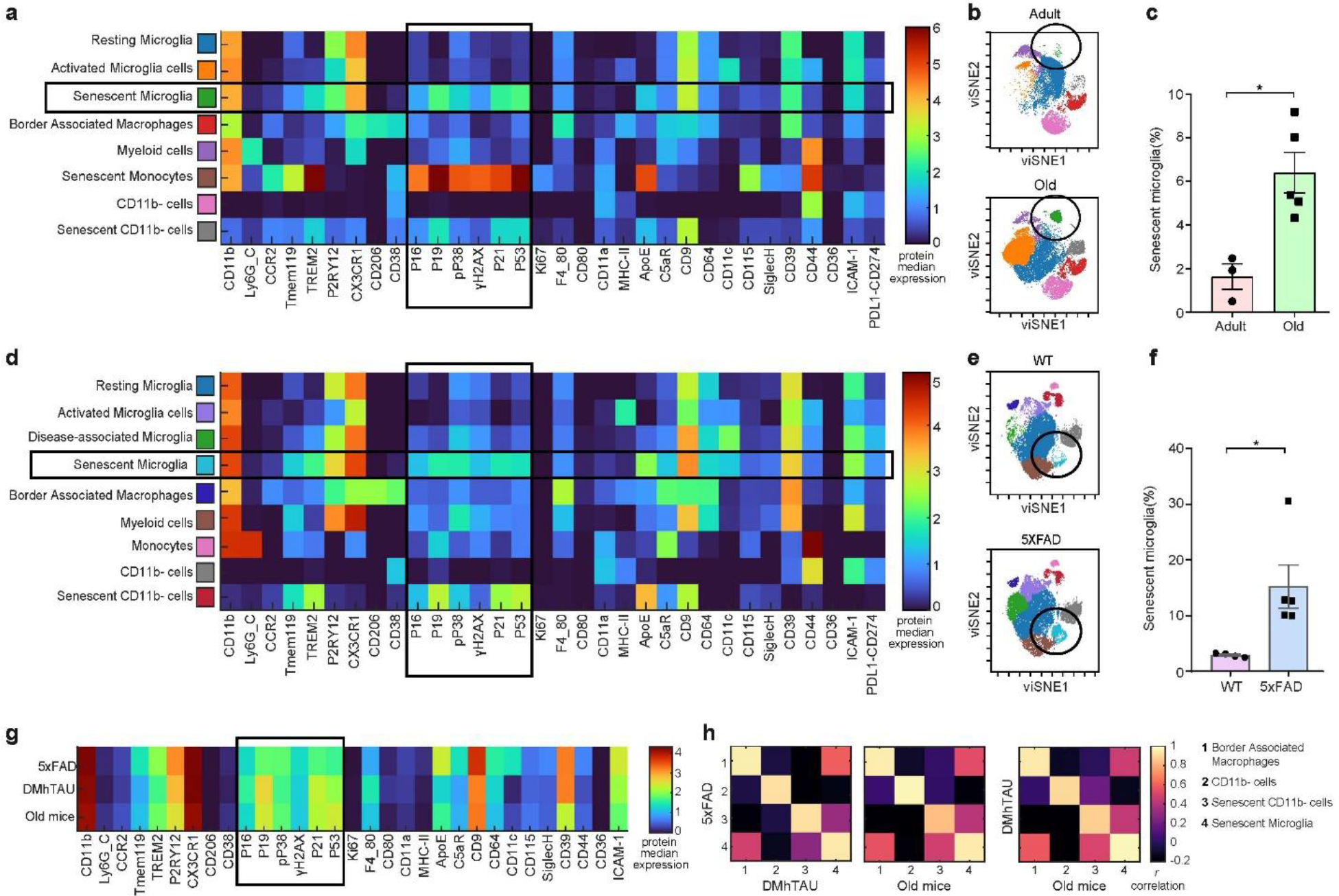
Senescent microglia with a distinct signature accumulate in aged mice and an AD mouse model. (a-c) 4-month-old C57BL/6 mice (Adult) (n = 3), and 24-month-old C57BL/6 mice (Old) (n = 5); (d-f) 11-month-old female mice (WT) (n = 4), and 11-month-old female 5xFAD mice (5xFAD) (n = 5). (a) Median marker expression values for each population. The presented matrix is the average profile across n=8 mice (b) Representative t-SNE plot of each experimental group. t-SNE map displaying 155,504 randomly sampled CD45+ cells, equally sampled. Colors correspond to FlowSOM-guided clustering of cell populations. (c) Quantitative analysis of the percentage of senescent microglia (as identified in a-b) in microglia (Student’s *t* test). (d) Median marker expression values for each population. The presented matrix is the average profile across n=9 mice. (e) Representative t-SNE plot of each experimental group. t-SNE map displaying 210,906 randomly sampled CD45+ cells, equally sampled. Colors correspond to FlowSOM-guided clustering of cell populations. (f) Quantitative analysis of the senescent microglia percentage in CNS microglia between the two experimental groups (Student’s *t* test). (g) Senescent microglia averaged Median marker expression values for each population. Each row represents a different experiment, first row is derived from 5xFAD mouse model of amyloidosis, the second from DMhTAU mouse model of tauopathy and third from aged mice. (h) Similarity in the expression profiles of four different cell-types across the three conditions, cluster 4 represent the senescent microglia. Data are presented as mean ± s.e.m.; *\*P < 0*.*05, **P < 0*.*01, ***P < 0*.*001*

### Senescent microglia are TREM2 dependent

To examine if TREM2 is an important player in the development of the senescent microglial phenotype, we compared levels of TREM2 in DAM and in senescent microglia, and found that within each tested mouse, levels of TREM2 expressed by senescent microglia were significantly higher than in DAM (Fig. 2a). Moreover, within the population of senescent microglia, TREM2 levels significantly correlated with the levels of senescence markers (Fig. 2b). We also assessed the levels of senescent microglia in *TREM2*^*-/-*^ 5xFAD mice, and found a significantly lower percentage of these cells, compared to *TREM*^*+/+*^5xFAD mice (Fig. 2c,d,e). Thus, although TREM2 is essential for microglial activation and for containing amyloid plaques^13,17,32^ its expression levels are found here to be associated with microglia transition towards senescence.

**Fig. 2.**
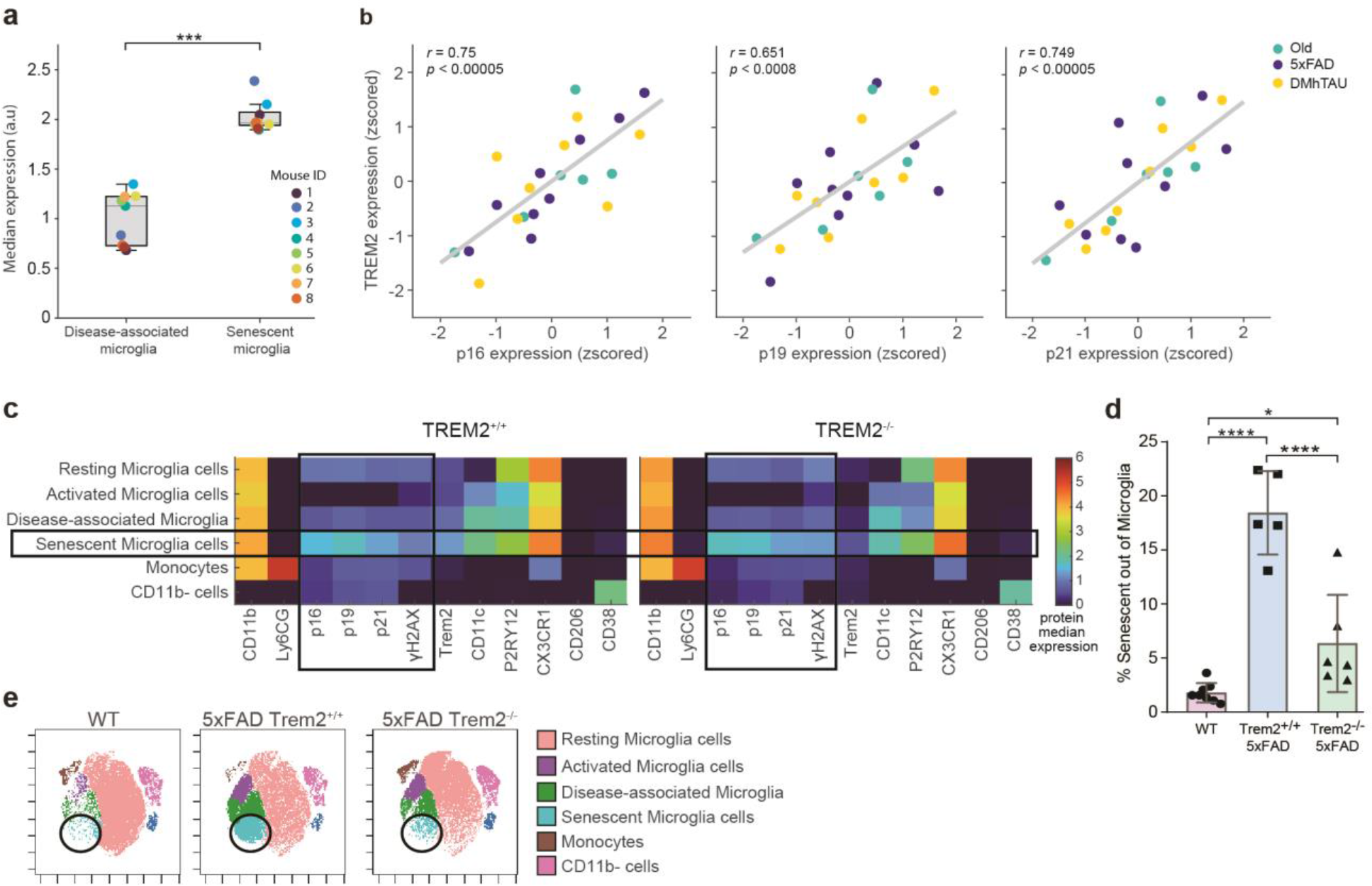
TREM2 is linked to senescent microglia accumulation. (a-e) 7-month-old female mice (WT) (n = 4), 7-month-old female mice TREM2^-/-^ (WT) (n = 3) compared to 7-month-old female *TREM*^*+/+*^5xFAD mice (n = 5), and 7-month-old female *TREM2*^*-/-*^5xFAD (n = 6). (a) Quantitative analysis of the median expression of TREM2 in each mouse between the clusters of DAM and of senescent microglia, each line represents one mouse (Student’s *t* test). (b) Person’s correlation of the expression of TREM2 with senescence markers, p16, p19 and p21. Each color represents a different experiment, the median expression was Z-scored to account for batch identity. (c) Median marker expression values for each population, one for each condition of *TREM2*^*+/+*^ and *TREM2*^*-/-*^. The presented matrix is the average profile across samples. (d) Quantitative analysis of the senescent microglia percentage within total CNS microglia between the three groups of each experiment (One-way ANOVA). (e) Representative t-SNE plot of each experimental group. The t-SNE map was created using 343,520 randomly sampled CD45+ cells, equally sampled from each brain, analyzed by mass cytometry. Colors correspond to FlowSOM-guided clustering of cell populations. Data are presented as mean ± s.e.m. One-way ANOVA was used for the analyses. *\*P < 0*.*05, **P < 0*.*01, ***P < 0*.*001*

### The transcript of senescent microglia

Since both activated microglia with the DAM signature^13^ and senescent microglia, found here, are TREM2 dependent, we further compared their profile. We found that senescent microglia appear distinct from DAM, as unlike DAM, they express high levels of homeostatic microglial markers (Extended Data Fig. 1). Next, we investigated their transcriptomic signature. We conducted a meta-analysis of previously published single nuclei RNA-sequencing data of WT, *TREM2*^*-/-*^WT, *TREM2*^*+/+*^5xFAD and *TREM2*^*-/-*^5xFAD mice^28^ (Fig 3a-c). We detected a cluster of senescent microglia, cluster 2, that appeared in 5xFAD mice but was almost entirely absent in the other groups (Fig. 3a,d-f). This cluster was identified as senescent using the SenMayo gene set^29^ (Fig. 3g). Cluster 2 showed a significant upregulation of TREM2, ApoE, CD9, and CD11c, similar to the senescent microglial cluster identified by CyTOF (Fig. 3h). Additionally, the transcriptional signature of cluster 2 was found to be similar to a previously described subtype of microglia, Highly-Activated Microglia, which only appear in old mice^30^ (Fig. 3h). Cluster 2 seemed to share many differentially-expressed genes (DEGs) with the Highly-Activated Microglia, including ApoE, Lpl, Lgals3, Cst7, Cd74, Cd63, Lilrb4a, Axl, Itgax, Cd83, Cd9, Lgals3bp, B2m, Csf2ra, Tyrobp, Crlf2, Cd34, Ccl4, Lyz2, Hif1a, Csf1, and Cd68^30^ (the list of all DEGs of each cluster is available in extended data Table 3).

**Fig. 3.**
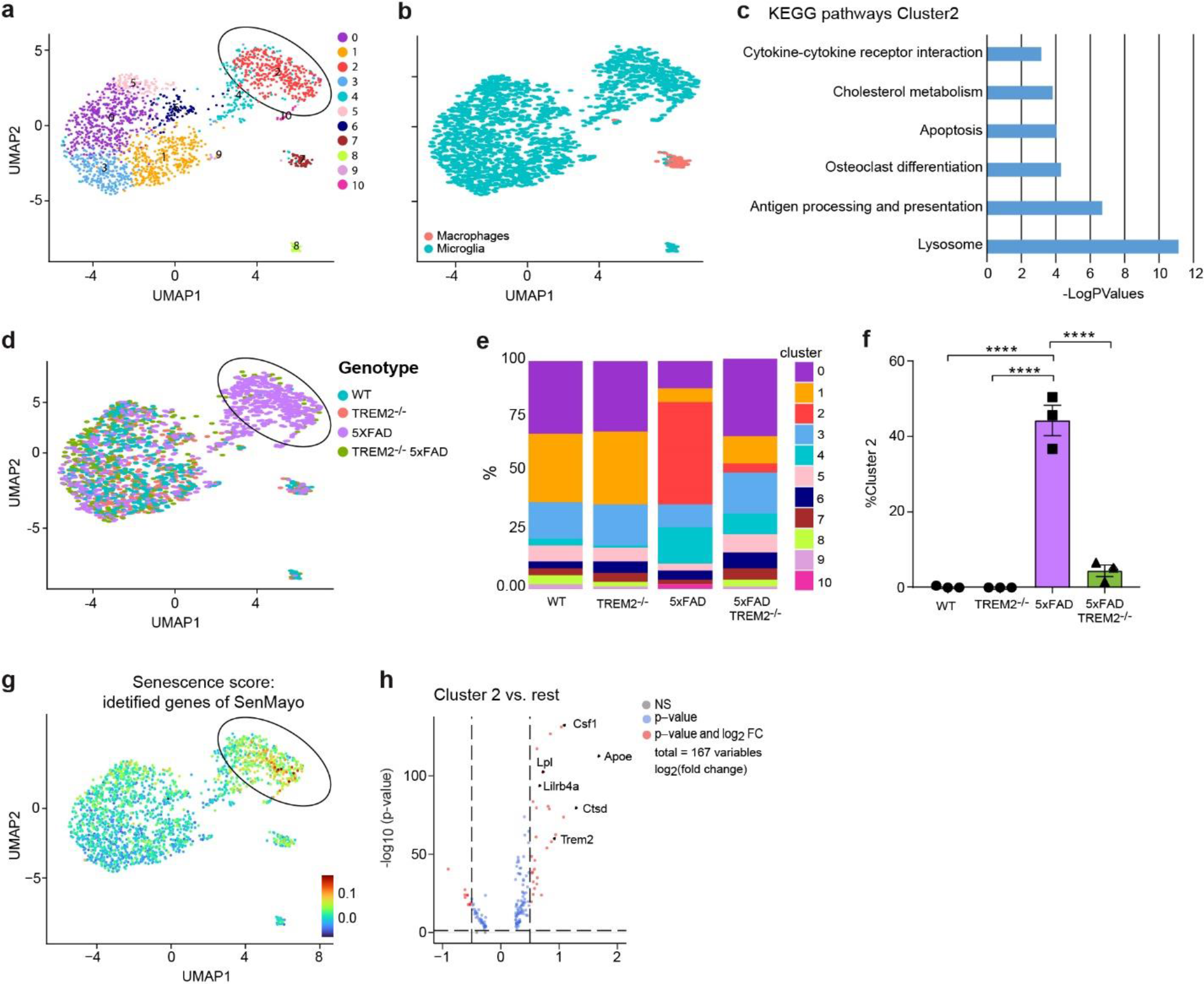
Transcriptional signature of senescent microglia. (a-c) Analysis of the single nuclei RNA-sequencing^28^ of 7-month-old female mice (WT) (n = 3), 7-month-old female *TREM2*^*-/-*^ mice (WT) (n = 3) compared to 7-month-old female TREM^+/+^5xFAD mice (n = 3), and 7-month-old female *TREM2*^*-/-*^5xFAD (n = 3)^28^. (a) Uniform manifold approximation and projection (UMAP) plots of re-clustered microglia and macrophages (b) UMAP plots of re-clustered microglia and macrophages coloured based of cell type (c) Pathways that were found significant in enrichment analysis (d) UMAP plots of re-clustered microglia and macrophages coloured based of genotype (e) The percentage of each cluster. (f) The percentage of cluster 2 across all groups (g) Module score of senescence signature based on SenMayo gene list for mice^29^, only 67 out of 120 were recognized in the data, identifying cluster 2 as enriched in senescent genes. (h) Volcano plot describing the results of differential analysis of genes Wilcoxon Rank Sum test – for the differential gene expression between cluster 2 and the rest of the clusters. Each dot represents one transcript. The horizontal line marks the significance threshold (Adjusted p-value, based on bonferroni correction using all genes in the data). The vertical dashed lines represent two- fold differences in expression.

### Senolysis of senescent microglia in AD

To examine the potential impact of TREM2-dependent senescent microglia on AD pathology, we treated 10-11 month-old 5XFAD mice with either 25mg/kg/day of the senolytic Bcl-2 family inhibitor, ABT-737^10,11^ or vehicle control. We first evaluated the effect of the treatment on recognition memory by using the Novel Object Recognition (NOR) test^31^, and subsequently on the accumulation of senescent microglia using CyTOF (Fig. 4a). We observed a significant improvement in memory, measured by NOR in the treated animals using two cohorts of mice (Fig 4b). In one cohort, following NOR, the brain was excised and further analyzed for levels of microglia by mass cytometry. The results revealed a significant reduction in the accumulation of senescent microglia in the treated mice (Fig. 4c-e). Senolytic treatment by ABT-737^10,11^ did not affect either the percentage of DAM (Extended Data Fig. 2), or the peripheral immune cell profile (Extended Data Fig. 3). Moreover, the level of TREM2-independent activated microglia was increased (Extended Data Fig. 2), further supporting the notion that TREM2-dependent senescent microglia are an outcome of a different activation trajectory compared to DAM. In the second cohort of mice, following treatment and the NOR task, we excised the mouse brains and performed quantitative real-time PCR to measure levels of genes encoding a panel of cytokines. We observed a significant reduction in levels of inflammatory cytokines in the 5XFAD mice treated with the senolytic drug relative to those treated with the vehicle control (Fig. 4f). This attributes the improved cognition observed in the 5xFAD mice to the depletion of senescent microglia and reduced inflammation. Altogether, senolysis, decreases the percentage of senescent microglia and neuro-inflammation, ameliorating the cognitive impairment in 5XFAD mice.

**Fig. 4.**
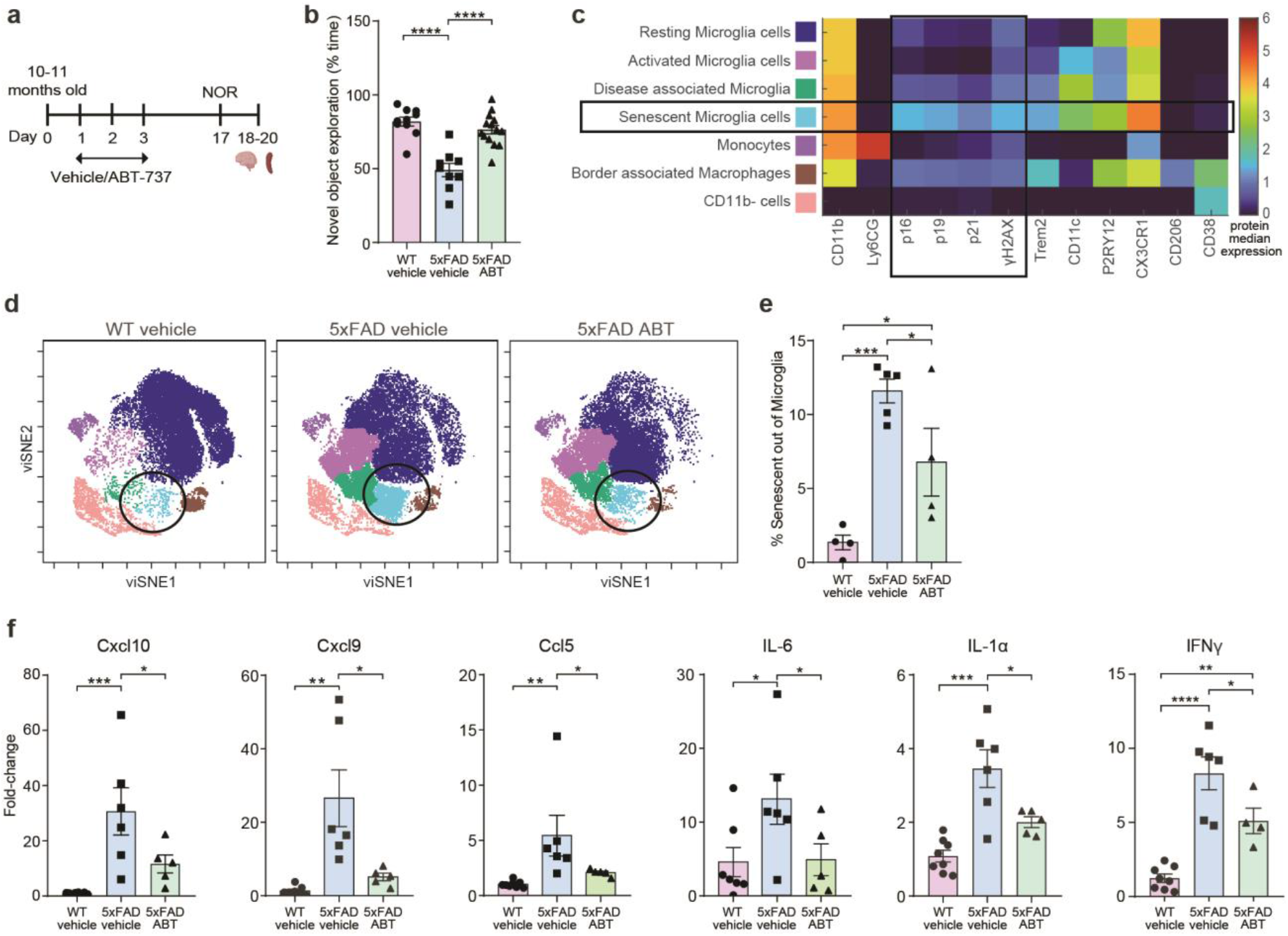
Senolytic therapy using ABT-737 ameliorates the accumulation of senescent microglia, cognitive decline and neuro-inflammation in 5XFAD mice. (a-f) First cohort of mice: 10-11-month-old mice injected with vehicle control (WT) (n = 4), compared to 10-11-month-old 5xFAD mice injected with vehicle control (n = 5), and 10-11-month-old 5xFAD mice injected with ABT-737 (n = 4). Second cohort of mice: 10-11-month-old mice injected with vehicle control (WT) (n = 5), compared to 10-11-month-old 5xFAD mice injected with vehicle control (n = 6), and 10-11-month-old 5xFAD mice injected with ABT-737 (n = 7). (a) Schematic representation of the experimental approach. (b) Novel Object Recognition (NOR): The memory recognition is presented as percent time the mouse interacted with the novel object divided by the total time spent with both objects. (c) Median marker expression values for each population. The presented matrix is the average profile across n=13 mice (d) Representative t-SNE plot of each experimental group. t-SNE map displaying 224,146 randomly sampled CD45+ cells, equally sampled from each sample, analyzed by mass cytometry. Colours correspond to FlowSOM-guided clustering of cell populations. (e) Quantitative analysis of the senescent microglia percentage in CNS microglia between the three groups of each experiment. (f) Quantitative PCR (qPCR) analysis. Data are presented as mean ± s.e.m. One-way ANOVA was used for the analyses. *\*P < 0*.*05, **P < 0*.*01, ***P < 0*.*001*

## Discussion

Here, we show that senescent microglia with a conserved signature accumulate in aging and in a mouse model of AD. We further show that this shared signature is characterized by high levels of TREM2 expression. Surprisingly, however, TREM2-dependent senescent microglia and the DAM population display distinct signatures, indicating that TREM2-activated microglia are not a homogenous population of cells. Eliminating TREM2-dependent senescent microglia using a senolytic drug, did not affect levels of microglia with a DAM signature, but led to an improvement in cognitive performance and to a reduction in the levels of genes encoding inflammatory cytokines in the brain.

Polymorphism of the TREM2 gene is associated with disease severity in AD patients^28^. Additionally, TREM2-expressing cells were reported to locate in close proximity to plaques^17,32^, and TREM2 facilitates microglial proliferation and their acquisition of the DAM profile^13,17^. It was suggested that TREM2 deficiency in animal models plays a role in AD progression^33,34^. Our data show that TREM2-dependent senescent microglia share markers, such as TMEM119, P2YR12, and CX3CR1, with homeostatic microglia. These results suggest that the two TREM2 expressing microglial populations, DAM and TREM2-dependent senescent microglia, do not represent a continuum, but are the outcome of distinct cellular states. This observation is in line with our findings here that the signature of senescent microglia is conserved across different conditions, including aging, AD and tauopathy. Our results could also explain why attempts to activate TREM2 as a therapeutic strategy have shown only limited beneficial effects in preclinical studies^35^. A recent study consistent with our findings, identified a microglial cluster that is upregulated in aged mice compared to juvenile mice, and expresses higher levels of TREM2^36^. In another study, a subtype of microglia with high expression of ApoE emerged following an injection of apoptotic neurons into WT mouse brains^37^. This microglial subtype was detected in aged and APP-PS1 mice, but was diminished in APP-PS1*TREM2*^*-/-*37^. Similarly, the cluster of senescent microglia also showed high expression of ApoE in both 5xFAD mice and aged mice, and was lower in *TREM2*^*-/-*^5xFAD mice. While the molecular mechanisms underlying the linkage of TREM2 to the senescent microglial phenotype are yet to be studied, our results highlight a strong connection between TREM2, microglial senescence, and AD. Other studies identified senescent microglia and astrocytes in mouse model of Tau pathology^38^. In another study, which did not use single-cell cytometry, senescent oligodendrocyte progenitor cells (OPCs) but not astrocytes or microglia, were identified in a mouse model of AD^39^. Of note, P16-positive microglia were also detected in aging^40^. Overall, results from different studies highlight the key role that senescent cells in the brain play in AD.

The senolytic drug, ABT-737^10,11^, which is a Bcl2 family inhibitor, improved cognitive performance in an AD mouse model. Although this was accompanied by the elimination of TREM2-dependent senescent microglia but not of DAM, we cannot rule out the possibility that the favorable outcomes are due to the elimination of additional senescent cells. In addition, the elimination of TREM2^+^ senescent microglia, with no significant effect on DAM, suggests that TREM2 displays a dual activity in microglia, which should be carefully considered when contemplating TREM2 as a therapeutic target^41^. Additionally, the improved cognitive performance in 5xFAD mice, following the elimination of TREM2-dependent senescent microglia was associated with a reduction in pro-inflammatory cytokines. This supports the contention that local brain inflammation is a major pathological factor that contributes to the cognitive deterioration found in human disease, as well^5^. This study suggests that targeting senescent cells may open new potential therapeutic strategies for AD, which might be effective even at late-disease stages. However, the senolytic drugs available today have limited applicability to humans due to their side effects^10,42^. In the future, developing new senolytic therapies that exclusively target senescent cells, or specifically, senescent microglia should be considered. In line with these contentions are the findings of the beneficial effect of targeting the inhibitory immune checkpoint pathway, PD-1/PD-L1 in animal models of AD and tauopathy, through a mechanism that involves reduced inflammation^43-46^ and anti-aging effect through a mechanism that reduces the abundance of senescent cells in all tissues tested^47,48^.

Overall, the present study highlights the role of senescent microglia in AD, and an unprecedented dual role for TREM2 in aging and dementia. While in the past, TREM2 was uniformly considered as a gene whose polymorphism is a risk factor in late-onset AD, here we show that despite its essential role, high levels of TREM2 could be detrimental. This complexity should be carefully considered when targeting TREM2 as a therapeutic approach.

## Methods

### Mice

Three mouse models were used in this study: (1) heterozygous 5xFAD transgenic mice (on a C57/BL6-SJL background), which express familial AD mutant forms of human APP (the Swedish mutation, K670N/M671L; the Florida mutation, I716V; and the London mutation, V717I) and PS1 (M146L/L286V) transgenes under transcriptional control of the neuron-specific mouse Ty-1 promoter^26^ (5xFAD line Tg6799; Te Jackson Laboratory); and (2) *Trem2*^*−/−*^5xFAD and *Trem2*^*+/+*^5xFAD mice on a C57/BL6 background were obtained from the laboratory of M. Colonna (Washington University), where they were generated as previously described^49^. (3) Heterozygous DM-hTAU transgenic mice, bearing two mutations (K257T/P301S) in the human-tau (hTAU) gene (double mutant, DM; on a BALBc-C57/BL6 background), associated with severe disease manifestations of frontotemporal-dementia in humans^27^, expressed under the natural tau promoter. Genotyping was performed by polymerase chain reaction (PCR) analysis of tail DNA. Throughout the study, WT controls in each experiment were non-transgene littermates from the relevant mouse colonies. Mice were bred and maintained by the animal breeding center of the Weizmann Institute of Science. All experiments detailed herein complied with the regulations formulated by the Institutional Animal Care and Use Committee of the Weizmann Institute of Science.

### Brain dissociation for single-cell suspension

Mice were euthanized using an overdose of ketamine–xylazine, followed by transcardial perfusion with cold PBS, and whole brains were excised. Next, the brains were cut using small seizures and incubated with RPMI supplemented with 0.4mg/ml Collagenase IV (Worthington Biochemical Corporation), 2mM HEPES (BI Industries), 10μg/ml DNase (Sigma), and 2% FCS]. Then gentle-MACS was used first brain program and incubated for 20min at 37°C shaker. Cells were homogenization again at the middle of the incubation, using the gentle-MACS brain program two after 10min. Finally, the sample went to gentle-MACS brain program 3 three times. The enzymatic reaction was stopped using cold RPMI supplemented with 0.5M EDTA (0.2%). Then, the cells were isolated using 30% Percoll (GE Healthcare, 17–0891–01) density gradient centrifugation (23,500g, 30min, 4°C). Then, cells were pelleted and rewashed.

### Antibody metal conjugation

Antibodies were conjugated to metals using the MIBItag Conjugation Kit (IONpath), according to the manufacturer’s protocols. Metals for conjugation were chosen to minimize noise and spillover between channels, according to guidelines in Han et al. (2018)^50^. Table S1 specifies the antibodies used and the metal allocated to each antibody.

### CyTOF sample preparation

After the brain single-cell extraction, the samples were stained by 1.25μM Cell-ID Cisplatin in MaxPar Cell Staining Buffer (Fluidigm). Then, the samples were washed twice with warmed RPMI supplemented with 10% FBS, followed by MaxPar Cell Staining Buffer. Next, they were incubated with Fc‐block CD16/32 (BD Biosciences; 10min, RT), followed by incubation with the extracellular antibodies (30min, RT). Then, the samples were fixed and resuspended with 4% Formaldehyde (Pierce) for 10min, washed, and kept on ice for 10 min. Next, they were permeabilized using 90% Methanol for 15min on ice, blocked again with 1% of Donkey serum, and stained with intracellular markers with 1% phosphatase inhibitor for 30 min. Lastly, the samples were washed twice and kept in 4% Formaldehyde (Pierce) with Iridium (125pM) at 4°C overnight. On the reading day, samples were washed twice with MaxPar Cell Staining Buffer, then washed twice with MaxPar water (Fluidigm) before the acquisition, using a CyTOF 2 Upgraded to Helios system (Fluidigm). A detailed list of antibodies used is supplied in the Supplementary section (Table S1). Purified antibodies were conjugated using MIBItag Conjugation Kit (IONpath).

### CyTOF Data processing

CyTOF data underwent the following pre-processing prior to analyses: First, the CyTOF software by Fluidigm was used to normalise and concatenate the acquired data. Then, several gates were applied using the Cytobank platform (Beckman Coulter): First, CD11b stable signal across time was gated, and then the event length and the Gaussian residual parameters. Then the beads were gated out using the 140Ce beads channel. Then, live single cells were gated using the cisplatin 195Pt, iridium DNA label in 193Ir, and single cells were gated using the two channels for iridium. Lastly, we gated for CD45-positive cells.

Gating strategy:

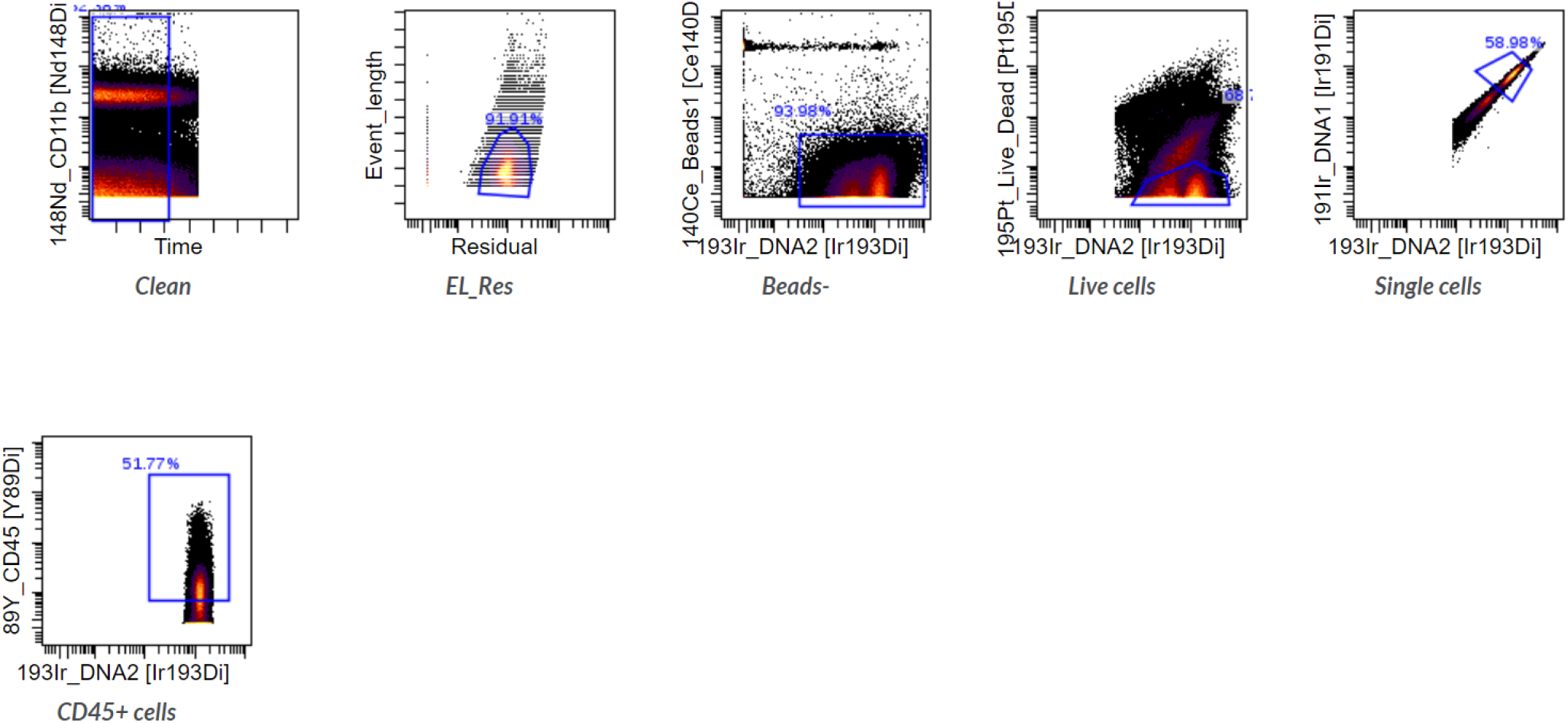

### CyTOF Data Analysis

CyTOF data were analyzed using Cytobank and Matlab. In Cytobank, CD45+ cells were further processed with equal sampling from each condition. The two-dimensional reduction was applied using the Cytobank “visne” (Vi-distributed Stochastic Neighbor Embedding; Vi-SNE) method, and FlowSOM-based clustering was performed. Using Matlab, the median expression of each cluster was averaged to create one Heatmap for each experiment. Additionally, t-tests and correlation analysis were done in Matlab. See raw data for the complete Matlab code.

### Cognitive assessment

Each mouse was subjected to a daily 3min handling session for five successive days prior to the first behavioural test. Behavioural studies were repeated twice, and results were combined. The investigators performing behavioural testing were blinded to the treatment group of the mice throughout the experiments. Testing sessions were recorded and analyzed using the EthoVision tracking system XT 11 (Noldus Information Technology).

The Behavioural results shown are consisted of two independent experiments. After completion of the behavior, the brain from one experiment were analyzed by CyTOF and the second experiment for levels of neuro-inflammation, by real-time PCR.

### Novel object recognition (NOR)

The novel object recognition test provides an index of recognition memory^31^. Modified from Bevins and Besheer^51^, a square grey box (45 × 45 × 50 cm) with visual cues on the walls was used. The task spanned two days and three trials: a habituation trial – a 20 min session in the empty apparatus (day 1). A familiarization trial – a 10 min session allowing the mice to interact with two identical objects (day 2), and a test trial: following a 1hr ITI, each mouse was returned to the apparatus for a 6 min session in which one of the objects was replaced by a novel one. Novel object preference was calculated as: per cent novel object exploration = ((novel object exploration time)/(novel object exploration time + familiar object exploration time)) × 100.

### RNA purification, cDNA synthesis and quantitative real-time PCR analysis

Mice were transcardially perfused with phosphate-buffered saline (PBS) before tissue excision. Cortex and hippocampus tissues were isolated from the brain under a dissecting microscope (Stemi DV4; Zeiss) and snap freeze in liquid nitrogen. Total RNA from them was extracted using the NucleoSpin RNA, Mini kit for RNA purification (Item number: 740955.50), and mRNA (∼2 μg) was converted into cDNA using a High Capacity cDNA Reverse Transcription Kit (Applied Biosystems). The expression of specific mRNAs was assayed using fluorescence-based quantitative real-time PCR (RT-qPCR) (Fast-SYBR PCR Master Mix; Applied Biosystems). Quantification reactions were performed in Duplicate for each sample using the ‘delta-delta Ct’ method. Hypoxanthine Phosphoribosyltransferase 1 (HPRT1) was chosen as a reference (housekeeping) gene. The following primers were used:

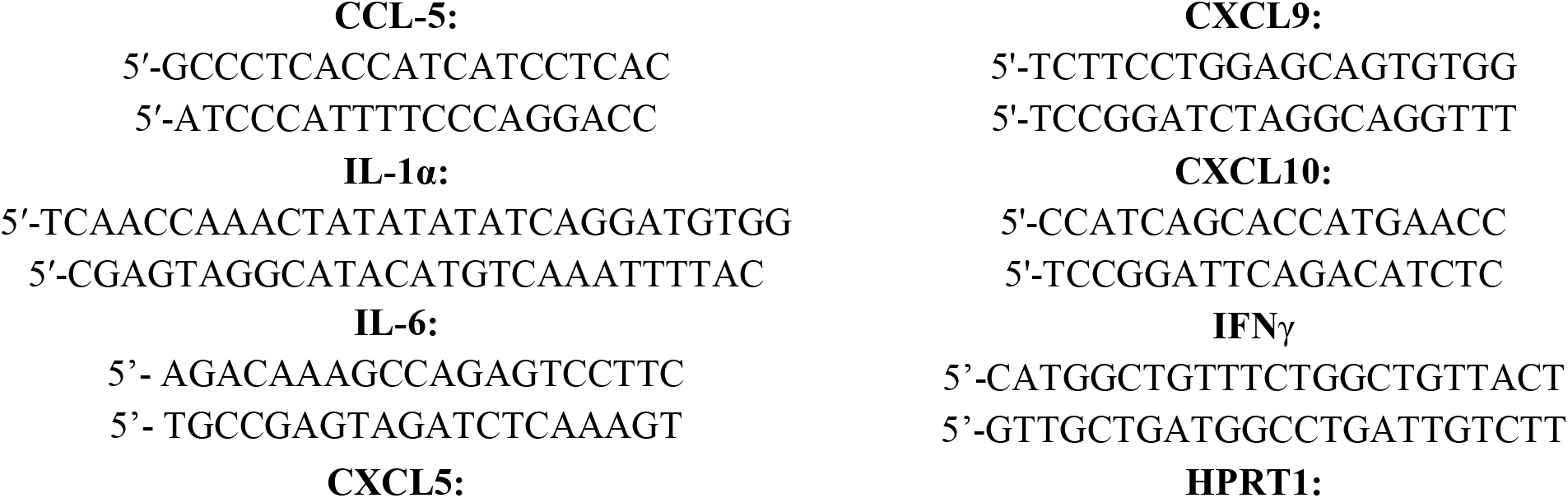
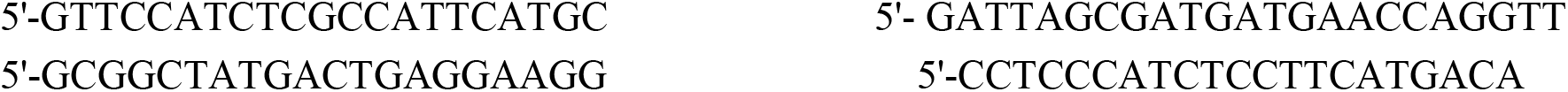

RT-qPCR reactions were performed and analyzed using StepOne software V2.2.2 (Applied Biosystems) and QuantStudio 3 software.

### Flow cytometry

Mice were transcardially perfused with PBS before tissue extraction. Spleens were mashed with the plunger of a syringe and treated with ammonium chloride potassium (ACK)-lysing buffer to remove erythrocytes. Before immunostaining, all samples were filtered through a 70-μm nylon mesh and blocked with anti-Fc CD16/32 (1:40; BD Biosciences). For FOXP3 staining, the samples were fixed, permeabilized, and subsequently stained using FOXP3/Transcription Factor Staining Buffer Set (eBioscience; 00–5523-00), according to the manufacturer’s instructions.

### Immunostaining panel

| **Abs Quantity**<br>(per 1x10<sup>6</sup> cells) | **CAT#** |
| --- | --- |
| Fc blocking (1:40) |  |
| PE-Cy7 αCD11b (1:200) | 101216 BioLegend |
| PerCP-Cy5.5 αCD45 (1:200) | 304028 BioLegend |
| PE-Dazzle αLy6C (1:200) | 128044 BioLegend |
| FITC αCD8a (1:200) | 100706 BioLegend |
| APC-Cy7 αTCRβ (1:200) | 109220 BioLegend |
| BV421 αCD4 (1:200) | 100544 BioLegend |
| Alexa 700 αCD44 (1:200) | 103026 BioLegend |
| PE αCD62L (1:200) | 104407 BioLegend |
| B220 BV650 (1:200) | 103241 BioLegend |
| Ly6-G BV605 1:200 | 127639 BioLegend |
| APC FOXP3 1:100 | 17-5773-82 eBioscience |
| Maxpar Cell Staining Buffer | Complete to 100ul per sample |

### Analysis of scRNA-sequencing data

We accessed the open access data from Zhou et al. (2020) (GSE140511). Details about sample processing, sequencing, and initial QC can be found in the original paper^28^. We accessed the gene-cell count matrices and cell barcodes data from 7-month-old female mice (WT) (n = 3), 7-month-old female mice *TREM2-/-* (n = 3), 7-month-old female *TREM+/+*5xFAD mice (n = 3), and 7-month-old female *TREM2-/-*5xFAD (n = 3). Analyses were performed using RStudio 2022.12.0. Following Seurat’s (v3.2.2) guidelines for quality control procedures, and under considerations to Zhou’s fine-tuning parameters used in their data analyses, we created a Seurat object, and negotiated data and statistical analyses. Briefly, cells having less than 200 genes and genes that appear in less than 5 cells were filtered out. Similarly, cells having more than 5% mitochondrial genes were also filtered out. Then, NormalizeData, ScaleData, RunPCA, FindNeighbors, FindClusters, and RunUMAP functions were activated according to object-tailored dimensions for Principal Components (PCs were estimated based on ‘JackStraw’ analysis and ElbowPlot). Cells type was determined using unsupervised mapping using ChenBrainData dataset from scRNAseq package and MouseRNAseqData dataset from celldex package. Afterwards, Microglia and macrophages cell types were subsetted, and Seurats’ objects were integrated to create one integrated object of microglial-macrophages cells type, on which we reperformed data dimensionality determination and scaling using SCTransform, RunPCA, FindNeighbors, FindClusters, and RunUMAP functions.

Clustering was performed selecting 8 principal components after ‘Jackstraw’ and ElbowPlot analyses, after which ‘clustree’ guided our selection for a resolution of 0.4 in FindClusters. Accordingly, RunUMAP was established.

Figures 3. a, b, d, e, g, h were created using DimPlot, FeaturePlot, ggplot2, and EnhancedVolcano packages. Figure 3.c is based on differentially expressed genes found between cells of cluster 2 and the rest of the cells in the object. Pathways were deducted from KEGG database using CytoScape V3.9.1, and STRING application. Significant pathways having Qvalues <0.05 are described in Excel column graph.

### Statistical analysis

The data was analyzed using a two-tailed Student’s t-test to compare between two groups, one-way ANOVA was used to compare several groups, we used either a Fisher’s LSD test for follow-up pairwise comparison, or Tukey correction, or Bonferroni correction, based on the different techniques. Data from behavioral tests were analyzed using two-way repeated-measures ANOVA and a Fisher’s LSD test for pairwise follow-up comparison. Sample sizes were chosen with adequate statistical power based on the literature and past experience, and mice were allocated to experimental groups according to age, gender and genotype. Investigators were blinded to the identity of the groups during experiments and outcome assessment. All inclusion and exclusion criteria were pre-established according to the IACUC guidelines. Results are presented as mean±s.e.m. In the graphs, y-axis error bars represent s.e.m. Statistical calculations were performed using GraphPad Prism software (GraphPad Software, San Diego, CA).

## Supporting information

ExData Table 3

## Acknowledgments

We sincerely thank Tsitsou-Kampeli A., Ben-Yehuda H., Suzzi S., Castellani G., Roitman L., Gal H. and Papismadov N. for technical assistance. We also thank all members of the Krizhanovsky laboratory and the Schwartz laboratory for helpful discussions. This study was supported by MS was supported by grants from the Advanced European Research Council grants 232835 and 741744, the European Seventh Framework Program HEALTH-2011 (279017), the Israel Science Foundation (ISF)-research grant no. 991/16, the ISF-Legacy Heritage Bio-medical Science Partnership research grant no. 1354/15, and the Thompson Foundation and Adelis Foundation (given to M. Schwartz); VK was supported by grants from the European Research Council H2020 program (856487), Centre for Research on Positive Neuroscience, Israel Science Foundation (1626/20), DFG (CRC 1506), Israel Ministry of Health, Belle S. and Irving E. Meller Center for the Biology of Aging, and Sagol Institute for Longevity Research. VK is an incumbent of The Georg F. Duckwitz Professorial Chair and Shimon and Golde Picker – Weizmann Award.

## Contributions

NR, MS and VK conceptualized the project. NR, MS and VK planned the experiments. NR designed and performed the CyTOF and Flow Cytometry experiments, and mice operations. SM performed and analysed the NOR task. UC and NR analysed Flow Cytometry. UC and NR injected mice. HA analysed the single-nucleus RNA data. NR and DD analysed the CyTOF results. HA, DD and NR performed data analysis. DE and NR performed and analysed RT-PCR. NR, TC and TMS established the CyTOF panel. JMPR established the Flow Cytometry panel. SM and UC assisted in experiments and experiment design. LC organized the mice colonies and performed genotyping. NR, MS and VK wrote the manuscript. All authors read and approved the final manuscript.

## Conflict of interest

V.K. is a co-inventor of patents on senolytics and senolytic approaches and consultant for Sentaur Bio. None of these influenced the data presented in this manuscript.

**Extended Data Fig. 1.**
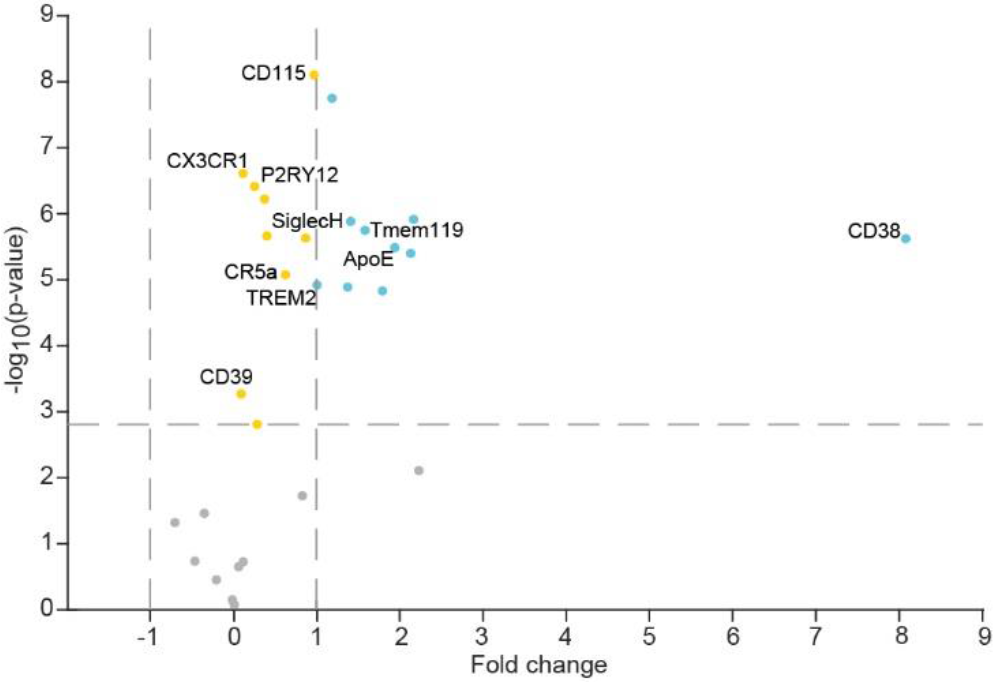
the differential protein expression between Disease-associated microglia and senescent microglia. Differentially expressed genes between Disease-associated microglia and senescent microglia. Each dot represents one protein. The horizontal line marks the significance threshold (P < 0.0016 after Bonferroni correction). The vertical dashed lines represent two fold differences in expression.

**Extended Data Fig Figure. 2.**
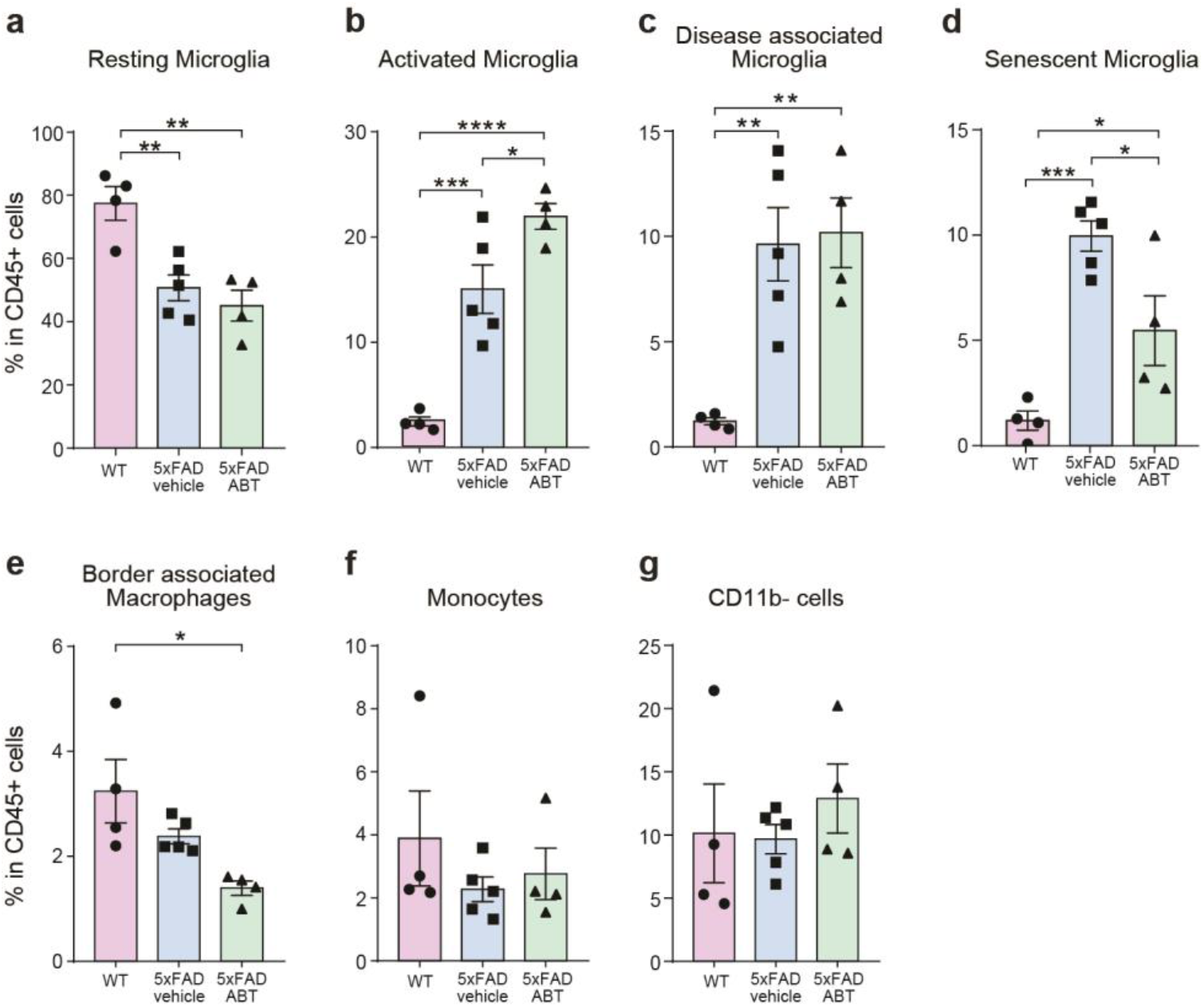
Other clusters percentages in CD45 cells. Quantitative analysis of the (a) Resting microglia, (b) activated microglia, (c) Disease associated microglia, (d) Senescent microglia, (e) Border-associated macrophages, (f) Monocytes and (g) CD11b-cells percentages in CNS CD45+ cells.

**Extended Data Fig. 3.**
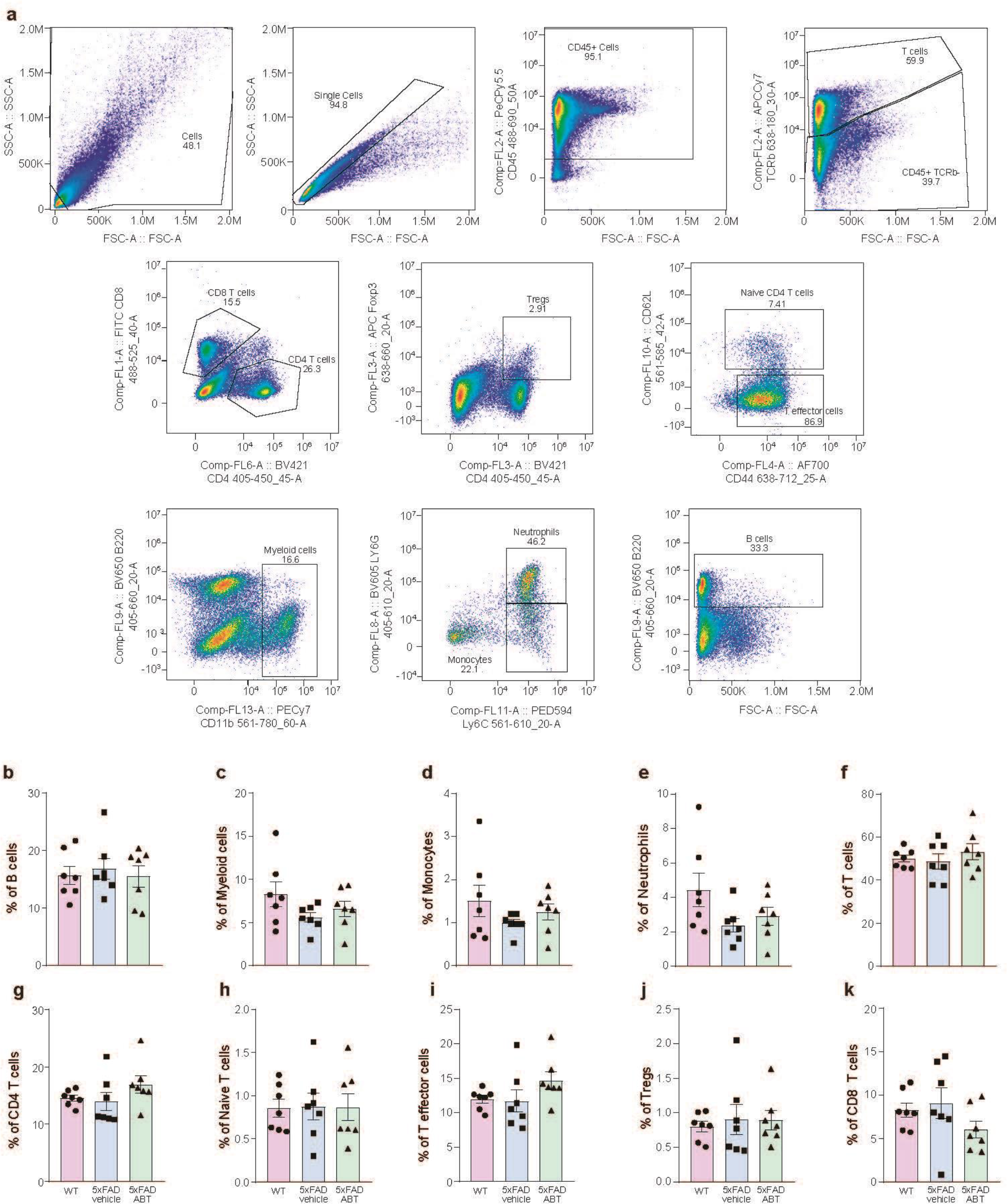
Flow-cytometry of splenocytes following ABT-737 treatment. (a) Gaiting strategy. (b-k) pink and circles represent the WT group, blue and squares represent the 5xFAD vehicle group, green and triangle represent 5xFAD ABT group (b) Quantitative analysis of B-cells percentage in total CD45+ cells. (c) Quantitative analysis of Myeloid cells percentage in total CD45+ cells. (d) Quantitative analysis of monocytes percentage in total CD45+ cells. (e) Quantitative analysis of neutrophils percentage in total CD45+ cells. (f) Quantitative analysis of T-cells percentage in total CD45+ cells. (g) Quantitative analysis of CD4+ T-cells percentage in total CD45+ cells. (h) Quantitative analysis of naïve T-cells percentage in total CD45+ cells. (i) Quantitative analysis of T-effector cells percentage in total CD45+ cells. (j) Quantitative analysis of T-regulatory cells percentage in total CD45+ cells. (k) Quantitative analysis of CD8+ T-cells percentage in total CD45+ cells.

**Extended Data Table 1.** CyTOF panel

| Marker | Metal | CAT# |
| --- | --- | --- |
| CD45 | 89Y | 3089005B |
| Ly6G/C | 141Pr | 3141005B |
| CD39 | 142Nd | 3142005B |
| p53 | 143Nd | BD554157 |
| CD115 | 144Nd | 3144012B |
| CD11a | 145Nd | BioLegend 153102 |
| F4/80 | 146Nd | 3146008B |
| CD36 | 147Sm | 3147018B |
| CD11b | 148Nd | 3148003B |
| γH2AX | 149Sm | NB100-384 |
| CD44 | 150Nd | 3150018B |
| CD64 | 151Eu | 3151012B |
| TERM2 | 152Sm | MAB17291 |
| PDL1-CD274 | 153Eu | 3153016B |
| CD38 | 154Sm | BioLegend 102702 |
| p21 | 155Gd | BD556431 |
| Siglec H | 156Gd | BioLegend 129602 |
| Tmem119 | 157Gd | AB-ab220249 |
| CD9 | 158Gd | 3158009B |
| P2RY12 | 160Gd | BLG-848002 |
| CD11c | 162Dy | 3162017B |
| ICAM-1 | 163Dy | 3163020B |
| CX3CR1 | 164Dy | 3164023B |
| p16 | 165Ho | AB-ab54210 |
| p19 | 166Er | AB-ab80 |
| CCR2 | 167Er | MAB55381R |
| Ki-67 | 168Er | 3168007B |
| CD206 | 169Tm | 3169021B |
| CD80 | 171Yb | 3171008B |
| ApoE | 173Yb | NB110-60531 |
| C5aR | 174Yb | BioLegend 135802 |
| pP38 | 176Yb | BLG-690202 |
| MHC-II (I-A/I-E) | 209Bi | 3209006B |

**Extended Data Table 2.** CyTOF small panel

| Marker | Metal | CAT# |
| --- | --- | --- |
| CD45 | 89Y | 3089005B |
| Ly6G/C | 141Pr | 3141005B |
| CD11b | 148Nd | 3148003B |
| γH2AX | 149Sm | NB100-384 |
| TERM2 | 152Sm | MAB17291 |
| CD38 | 154Sm | BioLegend 102702 |
| p21 | 155Gd | BD556431 |
| P2RY12 | 160Gd | BLG-848002 |
| CD11c | 162Dy | 3162017B |
| CX3CR1 | 164Dy | 3164023B |
| p16 | 165Ho | AB-ab54210 |
| p19 | 166Er | AB-ab80 |
| CD206 | 169Tm | 3169021B |

